# Fecal microbiota dysbiosis in macaques and humans within a shared environment

**DOI:** 10.1101/510065

**Authors:** Erica Grant, Randall C. Kyes, Pensri Kyes, Pauline Trinh, Vickie Ramirez, Tawatchai Tanee, Porntip Pinlaor, Rungtiwa Dangtakot, Peter M. Rabinowitz

## Abstract

Traditional zoonotic disease research focuses on detection of recognized pathogens and may miss opportunities to understand broader microbial transmission dynamics between humans, animals, and the environment. We studied human-macaque microbiome overlap in Kosum Phisai District, Maha Sarakham Province, Thailand, where a growing population of long-tailed macaques (*Macaca fascicularis*) in Kosumpee Forest Park interact with humans from an adjacent village. We surveyed workers in or near the park with elevated exposure to macaques to characterize tasks resulting in exposure to macaque feces in addition to dietary and lifestyle factors that influence gut microbiome composition. Fecal samples were collected from 12 exposed workers and 6 controls without macaque exposure, as well as 8 macaques from Kosumpee Forest Park and 4 from an isolated forest patch with minimal human contact. The V4 region of the 16S rRNA gene from fecal sample extracted DNA was amplified and sequenced using Illumina HiSeq to characterize the microbial community. A permuted betadisper test on the weighted UniFrac distances revealed significant differences in the dispersion patterns of gut microbiota from exposed and control macaques (p=0.03). The high variance in gut microbiota composition of macaques in contact with humans has potential implications for gut microbiome stability and susceptibility to disease, described by the Anna Karenina principle (AKP). Human samples had homogenous variance in beta diversity but different spatial medians between groups (p=0.02), indicating a shift in microbial composition that may be explained by fundamental lifestyle differences between the groups unrelated to exposure status. SourceTracker was used to estimate the percent of gut taxa in exposed humans that was contributed by macaques. While one worker showed evidence of elevated contribution, the overall trend was not significant. Task observations among workers revealed opportunities to employ protective measures or training to reduce exposure to occupational hazards. These results suggest the potential for hygiene measures to mitigate negative aspects of contact between humans and macaques in order to optimize the health of both populations.

## Introduction

### Background

Habitat fragmentation and human encroachment results in a patchwork of isolated non-human primate populations across Thailand with potential for increased human-macaque contact [1]. Supplemental feeding for religious reasons or tourism contributes to a growing macaque population unconstrained by natural food resources. Amidst these changes, the high level of human-macaque conflict has led researchers to call for improved management plans and conservation strategies [1, 2]. The reasons for concern are two-fold: 1) human-macaque conflict—such as crop-raiding—can disrupt or damage livelihoods, resulting in negative perception of macaques and impairing conservation efforts and 2) the increased level of contact can provide opportunities for transmission of zoonotic diseases into either macaque or human populations.

Due to their genetic similarity, humans and macaques are susceptible to many of the same infectious diseases including tuberculosis and hepatitis [3]. Parasitic infections from soil-transmitted helminths capable of infecting humans and macaques, such as *Strongyloides fuelleborni* and *S. stercoralis* have also been documented in this region [4, 5]. These pathogens may (e.g. Ebola virus) or may not (e.g. Herpes B virus) cause overt disease in macaques, however cross-species pathogen transmission from wildlife reservoirs to human hosts is a recognized factor in the emergence of novel diseases [6]. These spillover events can readily occur where natural and urban spaces meet. An example of such an interface is the Kosumpee Forest Park (KFP), a small fragmented forest in northeastern Thailand that is home to over 700 long-tailed macaques (*Macaca fascicularis*) and lies adjacent to the Kosum Phisai community of 4,235 persons [7, 8]. Unlike other popular tourist sites in Southeast Asia, macaques in KFP rarely climb onto people and feeding by the people often involves simply throwing food on the ground [9]. However, workers in and around the park regularly feed macaques and sweep macaque excrement from public spaces, elevating their risk of zoonotic disease transmission relative to other members of the Kosum Phisai community. While these workers represent a control point for broader spread of zoonoses, little is known about their knowledge, attitudes, and practices surrounding macaque exposure [10]. Based on task observations, it may be possible to identify intervention strategies to reduce exposure to macaque biological material. Such strategies could include the use of personal protective equipment (PPE), training in basic hand hygiene, or other measures to mitigate the risk of disease transmission. These measures would also promote responsible wildlife conservation by protecting macaques from pathogens that the workers could transmit through reverse zoonotic transmission.

The zoonotic spillover potential of certain pathogens, such as simian foamy virus, can be investigated through blood sample collection, however this method can be logistically challenging since it requires trapping and immobilization of wild macaques. Additionally, previous surveys among this worker population indicated that scratches or bites are infrequent, and ingestion of aerosolized fecal matter may be a more common route of exposure to zoonoses. Therefore, in this setting, we chose to analyze the microbial communities of fecal samples in humans and macaques with close contact. Advantages of analyzing the fecal microbiota include the fact that, compared to the skin microbiota, it is better characterized in literature, more temporally stable, and yield higher read counts [11]. Recent studies have demonstrated that the community composition of human microbiota is influenced by our environment and the animals sharing that environment. The degree of contribution from these sources can be quantified using Bayesian approaches like SourceTracker [12, 13]. We performed a pilot study of fecal microbiota of workers and macaques in a shared environment to test the hypothesis that workers exposed to macaques will exhibit microbiota profiles that contain a greater percentage of microbes found in macaque feces compared to unexposed individuals. The goals of this study were to provide a baseline assessment of the risk of zoonotic disease transmission between macaques and workers and guide prevention recommendations.

## Materials and methods

### Study design

This pilot study was a cross-sectional sampling of humans and macaques, comparing humans with occupational contact with macaques (exposed humans) to humans without such contact (human controls), and macaques in close contact with humans (exposed macaques) compared to macaques without significant human contact (control macaques).

### Human participants

Eligible workers (n=12) were defined as members of the community who contact macaques or macaque bodily fluids (blood, feces, urine) as a component of their paid work at least once per week. Workers were excluded if they had not worked at that site for a minimum of three months. Human controls (n=6) were recruited from a convenience sampling of adults at Mahasarakham University and were eligible if they were over 18 years of age and reported no contact with macaques. Recruited participants were informed of study objectives and their rights as participants and offered 100 Thai baht as compensation for their time.

### Animal participants

Exposed macaques (n=8) were sampled at Kosumpee Forest Park, with an effort to collect samples from macaques belonging to each of the social groups and age/sex distribution representative of the overall population. These macaques are individually identifiable by facial features or other unique characteristics by RCK. Control macaques (n=4) were sampled from a nearby forest in Phon Ngam in the same manner as exposed macaques, and age/sex were recorded.

### Environment

Sites were selected based on the level of interaction between humans and macaques. The study site for exposed human participants was a village of approximately 4,235 individuals adjacent to Kosumpee Forest Park (KFP), Kosum Phisai District, Maha Sarakham Province in northeastern Thailand (16°15'19"N 103°04'06"E) [8]. The forest park is an isolated forest patch of approximately 0.2 km^2^, bordered on the east by the Chi River and to the south by the Kosum Phisai village. The park contains over 700 long-tailed macaques, divided into five social groups with largely overlapping ranges [7]. Control sites were Mahasarakham University for humans, approximately 24 km E of KFP, and a small forest tract in Phon Ngam (16°21’01“N 102°56’54”E) for macaques, approximately 16 km NW of KFP, where there is minimal human-macaque interaction.

### Measurement

Interviews, task observation, and sample collection was conducted from Sept 24 – Oct 7, 2017. Survey data and sample metadata were collected and stored using the REDCap electronic database [14].

Macaque workers were surveyed regarding practices, training (e.g. macaque behavior, PPE use, wound care) and their knowledge of the principle that macaques and humans can share diseases. We piloted the occupational risk factor survey used in this study for eight park workers in October 2016 and revised it to address limitations that emerged during administration and analysis. Additions included a dietary questionnaire based on a modified food frequency questionnaire (FFQ). The full occupational questionnaire is available in supplemental materials. Task observations of workers were recorded using a GoPro HERO5 video recorder (GoPro, Inc., San Mateo, CA, USA) in order to assess work activities and supplement characterization of exposure opportunities identified in the survey. Task observations were performed at the job title level (vendor, park worker, and janitor), not for each individual, for feasibility. The scoring criteria was devised by authors based on probable routes for fecal microbe transmission to humans. The video recordings were reviewed by two individuals to maintain consistency and discrepancies were addressed by reexamining the video segment. During review of recorded tasks, an exposure category was assigned by the reviewer at 5 minute intervals, based on proximity of macaques (high=direct contact or within 3m, low=beyond 3m or not visible) and behavior (aerosol generation or hand-to-mouth contact).

Fresh fecal samples were placed immediately into OMNIgene.GUT kits (DNA Genotek, Ontario, Canada) to stabilize and preserve microbial community composition and stabilize DNA in the absence of a cold chain. Workers were provided with sterile collection kits and instructions in Issan Thai for proper specimen collection; macaque samples were similarly collected using a sterile spatula from the center of fresh excrement. Samples stored at ambient temperature as per OMNIgene.GUT kit instructions until they were processed at Khon Kaen University. QIAamp PowerFecal DNA Isolation kit (Qiagen, Hilden, Germany) was used to extract genomic DNA, following manufacturer protocols. DNA concentration was determined using a NanoDrop2000 spectrophotometer (NanoDrop Technologies Inc., DE, USA) and the integrity of DNA was evaluated by running 5 ul of sample on a 0.8% agarose gel under 100 V for 30 min and assessing bands. Extracted DNA samples were shipped overnight on blue ice to Genewiz Laboratories in Suzhou, China. DNA quality was verified by Genewiz using NanoDrop, Qubit, and agarose electrophoresis. The V4 region of the bacterial 16S genes were amplified using the 515F-806R primers, based on the Earth Microbiome Project protocol [15]. Amplicons were sequenced on an Illumina HiSeq platform by Genewiz Laboratories. Raw FASTQ files and metadata can be accessed through the Qiita database (https://qiita.ucsd.edu/) (accession no. 11835) and the European Bioinformatics Institute, European Nucleotide Archive (accession no. ERP111664).

### Analysis

DNA sequences or reads in the form of FASTQ files were analyzed with QIIME2 version 2017.12.0 pipeline [16]. DADA2 version 2017.12.1 was used for sequence quality control and feature table construction [17]. Forward reads were truncated to 280 bp and reverse reads to 260 bp. Alpha diversity metrics (observed OTUs, Shannon’s diversity index, Faith’s Phylogenetic Diversity, and Pielou’s Evenness) were calculated in QIIME2. In order to attain valid comparisons of abundance and diversity across samples, we normalized to the lowest sample depth of 12,466 reads per sample [18]. Sequences were assigned taxonomy using the SILVA 132 reference database [19]. Analysis of Composition of Microbiomes (ANCOM) was performed in QIIME2 between species and exposure groups with significantly different abundance values identified based on the W-statistic [20]. Principal Coordinate of Analysis (PCoA) plots and taxa bar plots were generated using the phyloseq package (version 1.22.3) in R [21]. PCoA plots were generated to visualize clustering patterns based on weighted UniFrac distance measures, which describes the degree of similarity between sample compositions by measuring the fraction of unique branch length from the phylogenetic tree of sample features and weights the distance by the relative abundance of that taxa within a sample. Profile clustering patterns from weighted UniFrac distance measures were analysed using adonis and betadisper tests from the vegan package (version 2.5.1) [22]. All tests were performed using 999 permutations based on the spatial median. To further characterize microbial sharing, SourceTracker [23] was applied to feature tables with macaques as source and humans as the sink under the default settings at a rarefaction depth of 1000 with 100 burn-ins and 10 re-starts.

### Study team

The COHERE guidelines for reporting of One Health studies were followed in the preparation of this manuscript [24]. Study members represented the following areas of expertise: primatology (RCK, PK), human health (PR), anthropological medicine (VR), microbial ecology (EG), molecular biology (PP, RD), computational biology (PT), and environment/resource management (TT).

### Ethics statement

The research in this study was approved through the University of Washington Institutional Review Board (IRB) for human subjects research and Institutional Animal Care and Use Committee (IACUC) for animal research (#51546 and #3143-04, respectively). The study also received approval through Mahasarakham University for human and animal subjects research (protocol numbers 037/2016 and 0009/2016, respectively). Written informed consent was obtained from all human participants and they were informed that participation was voluntary, they could withdraw at any time, and questionnaire responses, individual microbiota results, and task observation videos would be kept confidential and de-identified. Macaque samples were obtained from fresh defecations, therefore no direct macaque handling occurred as part of this study. This study was part of a larger project approved by the National Research Council of Thailand (NRCT project approval to RCK - Project ID: 2016/048; “Healthy Coexistence between Human and Non-human Primates: A One Health Approach”).

## Results

### Questionnaire

Exposed workers included government employees of Kosumpee Forest Park (n=8), janitors at a nearby school (n=3), and a vendor stationed near the park entrance (n=1). All study participants were born in Thailand and lived in the Maha Sarakham province for over a year. Demographic factors are summarized in Table 1.

**Table 1.** Human metadata.

| Factor | Exposed (n=12) | Control (n=6) |
| --- | --- | --- |
| Age, years (mean ± SD) | 47.17 ± 11.36 | 27.5 ± 9.44 |
| Sex |  |  |
| Male | 75% (9) | 50% (3) |
| Female | 25% (3) | 50% (3) |
| Education, years (mean ± SD) | 9.0 ± 3.05 | 16.8 ± 5.76 <sup>1</sup> |
| Household size |  |  |
| 1-3 | 25% (3) | 67% (4) |
| 4-6 | 58% (7) | 33% (2) |
| 7-9 | 17% (2) | 0 |
| Self-rated general health |  |  |
| Fair | 77% (8) | 0 |
| Good | 33% (4) | 83% (5) |
| Excellent | 0 | 17% (1) |
| Smoker | 75% (9) | 0 |
| Health problems in past year |  |  |
| Fever | 92% (11) | 67% (4) <sup>1</sup> |
| Respiratory problems | 58% (7) | 67% (4) <sup>1</sup> |
| Gastrointestinal problems | 33% (4) | 67% (4) |
| Skin problems | 25% (3) | 0 |
| Infectious diseases in lifetime |  |  |
| Tuberculosis | 8% (1) | 0 |
| Malaria | 8% (1) | 0 |
| Dengue | 17% (2) | 0 |
| Other parasites, hookworm | 58% (7) <sup>1</sup> | 0 <sup>1</sup> |
| Vaginal birth method | 77% (8) | 50% (3) |
| Breast-fed as infant | 92% (11) | 33% (2) |
| BMI | 25.5 ± 5.8 | 23.8 ± 3.5 |
| Antibiotic use in past month | 17% (2) <sup>4</sup> | 33% (2) <sup>2</sup> |

Demographic, early life history, dietary, and other health factors for exposed and control humans, which may influence gut microbiota or may be related to macaque exposure. Superscripted numbers reflect the number of missing datapoints.

Occupational factors related to microbial transmission are presented in Table 2. More than half of workers regularly wash hands without soap. All participants reported handwashing before and after eating (not listed in table), however task observation footage suggested this was not the case for at least four participants. PPE use as reported in the survey was low, which was further confirmed by the video recorded task observations. Respondents did not report receiving training relevant to safe animal handling or disease prevention before working around macaques. In an assessment of zoonotic disease knowledge, one-third of workers thought a diseased animal could transmit that agent to a human. Only one worker thought a human could make an animal sick and remarked that this would be with a high degree of contact. Workers typically only have direct physical contact with carcasses, but occasionally trap live macaques to move them from private properties to the forest park or when helping researchers. In one instance, a janitor had to remove a macaque from a classroom using a stick and grabbing it by hand. When around animals that appear sick, workers’ primary form of precaution was to avoid contact.

**Table 2.** Occupational risk factors.

| Factor | Response |
| --- | --- |
| Years at current job (mean ± SD) | 18.40 ± 11.79 |
| Hrs/wk around macaques or their feces (mean ± SD) | 45.08 ± 8.694 |
| Handwashing |  |
| Water only | 58% (7) |
| Soap and water | 50% (6) |
| Alcohol-based sanitizer | 8% (1) |
| PPE |  |
| Disposable gloves | 8% (1) |
| Paper or cloth dust masks | 17% (2) |
| Rubber boots | 25% (3) |
| Received animal/disease safety training <sup>a</sup> | 0% (0) |
| Change in macaque behavior | 42% (5) |
| Knowledge of animal to human transmission | 33% (4) |
| Concerned about diseases from animals at work | 33% (4) |
| Knowledge of human to animal transmission | 8% (1) |
| Take precautions around animals that look sick | 83% (10) |

Occupational risk factors related to macaque exposure among park workers, janitors and vendors based on a questionnaire.
<sup>a</sup> Training topics included animal behavior, animal capture/restraint, infectious disease prevention, PPE use, or wound care.

Since starting their current job, workers noted that macaques seem “naughtier”, wait for provisioning or do not look for natural food, and eat more human food (e.g. chicken, meatballs, soda). All workers reported finding macaques that looked sick or had died. Carcasses were typically buried or burned. One janitor remarked that, “Last month 3 monkeys die, pick them up by broom into plastic bag and then threw them into the forest.”

Workers were asked what diseases they were primarily concerned about getting in general, not necessarily from macaques. Responses included leptospirosis (n=3), cancer (n=2), the common cold (n=2), cirrhosis (n=1), allergies (n=1), and an airborne infectious disease (n=1) (Table 1). One worker was concerned about a “disease that come with monkey poo because I have to sweep it every day.” In contrast, non-communicable diseases like high blood pressure (n=3), cancer (n=1), diabetes (n=1) and hemorrhoids or constipation (n=1) were the primary disease concerns among controls.

An abbreviated food frequency questionnaire revealed dietary differences in the type of animal protein consumed. Control group members consumed more pork (p=0.04) and snail (p=0.03), whereas exposed workers typically ate more frog (p=0.04). There were no significant differences found in other dietary categories, including raw meat consumption. All respondents reported that they pass normal formed stool (Type 3/4 on Bristol stool scale), except one, from the exposed group, who reported Type 1/2.

### Task observation of workers

Park workers engaged in the highest exposure activities based on recorded task observations, followed by individuals working as school janitors, then vendors (Table 3). Using the number of exposure events divided time observed to calculate relative risk (RR), a park worker is 1.78 times more likely than a school janitor and 2.84 times more likely than a vendor to work within 3m of macaques or engage in risk elevating activities (e.g. aerosol generation, hand-mouth contact) during the task observation.

**Table 3.** Potential exposure to macaque feces based on video-recorded task observation

| Occupation | Proximity to macaques or macaque feces |  |  |  | Risk elevation<sup>c</sup> | Time observed | RR (95% CI) |
| --- | --- | --- | --- | --- | --- | --- | --- |
|  | Not visible<sup>a</sup> | Beyond 3m<sup>a</sup> | Within 3m<sup>b</sup> | Contact<sup>b</sup> |  |  |  |
| Vendor | 0 | 16 | 3 | 0 | 4 | 95 min | Ref |
| Janitor | 4 | 7 | 2 | 0 | 11 | 65 min | 1.78 (1.67-1.90) |
| Park worker | 0 | 3 | 6 | 4 | 9 | 65 min | 2.84 (2.67-3.02) |

Task observation assessment for the three worker groups (vendor, janitor, and park worker) reveals variation in proximity to macaques and behaviors that elevate risk of potential exposure. Risk of a potential exposure event during the task observation is presented relative to the lowest exposure occupation (vendor).
<sup>a</sup> No exposure
<sup>b</sup> Exposure
<sup>c</sup> Exposure, defined as engagement in activities that generate aerosols and/or eating, drinking, or smoking without prior handwashing

### Macaque demographics

Macaques were age and sex identified according to Table 4. Among exposed macaques, members of four of the five social groups within Kosumpee Forest Park are represented in this study. Two individuals were sampled from each group (i.e. Red Dot, Stump Tail, Droop Lip and Hare Lip). At the control site, all four members were from the same social group.

**Table 4.**
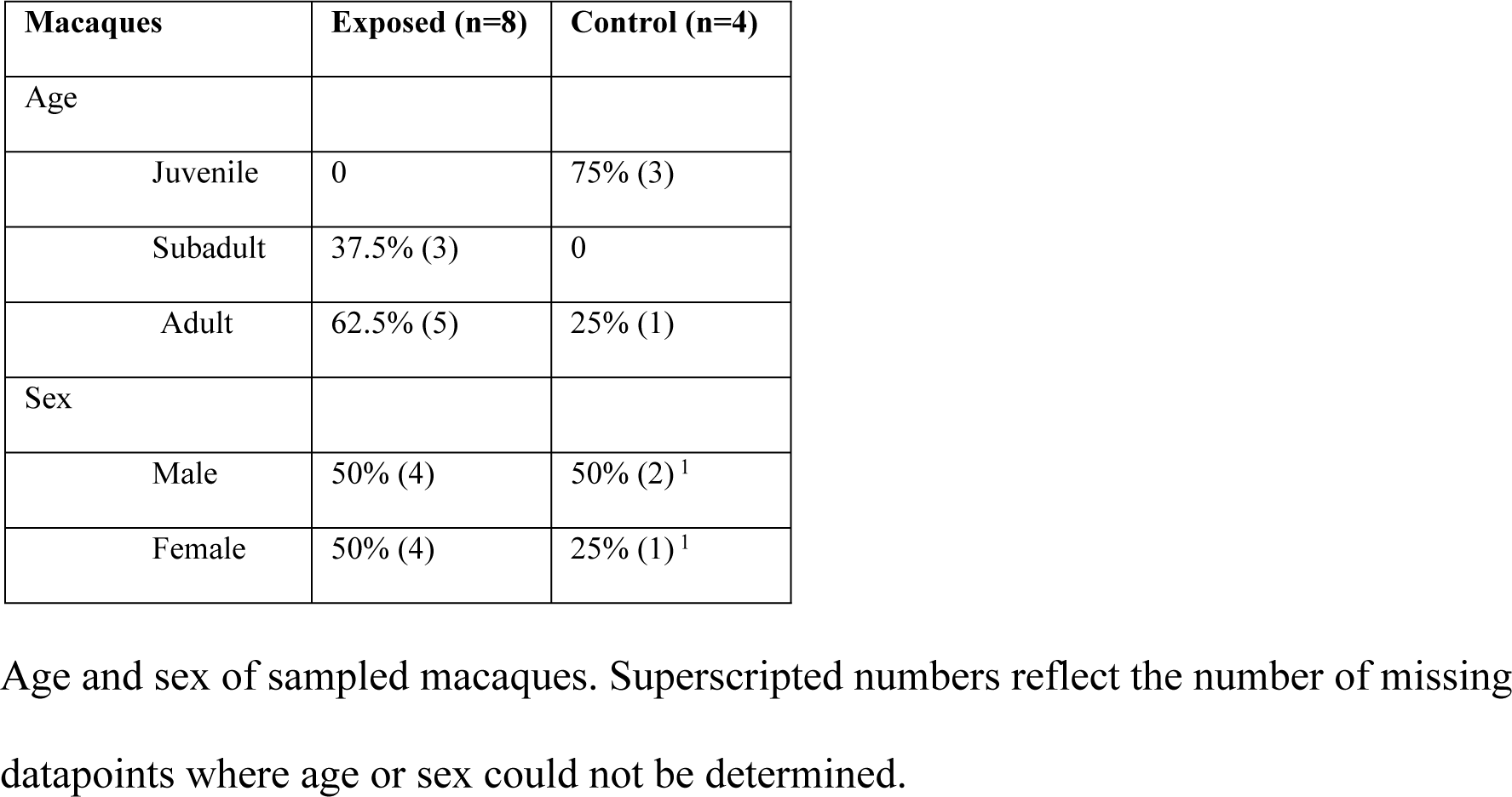
Macaque metadata.

| Macaques | Exposed (n=8) | Control (n=4) |
| --- | --- | --- |
| Age |  |  |
| Juvenile | 0 | 75% (3) |
| Subadult | 37.5% (3) | 0 |
| Adult | 62.5% (5) | 25% (1) |
| Sex |  |  |
| Male | 50% (4) | 50% (2) <sup>1</sup> |
| Female | 50% (4) | 25% (1) <sup>1</sup> |

Age and sex of sampled macaques. Superscripted numbers reflect the number of missing datapoints where age or sex could not be determined.

## Fecal microbiota analysis

### Phylum-level abundance

A total of 3,307 amplicon sequence variants (ASVs) were generated from 628,623 total read counts. There was an average of 20,954 reads per sample (range: 12,466-35,318). Fig. 1 shows the relative abundance of bacterial phyla in each sample, after rarefication to minimum sample size. All sample profiles were dominated by Bacteroidetes, Firmicutes, and Proteobacteria.

**Figure 1.**
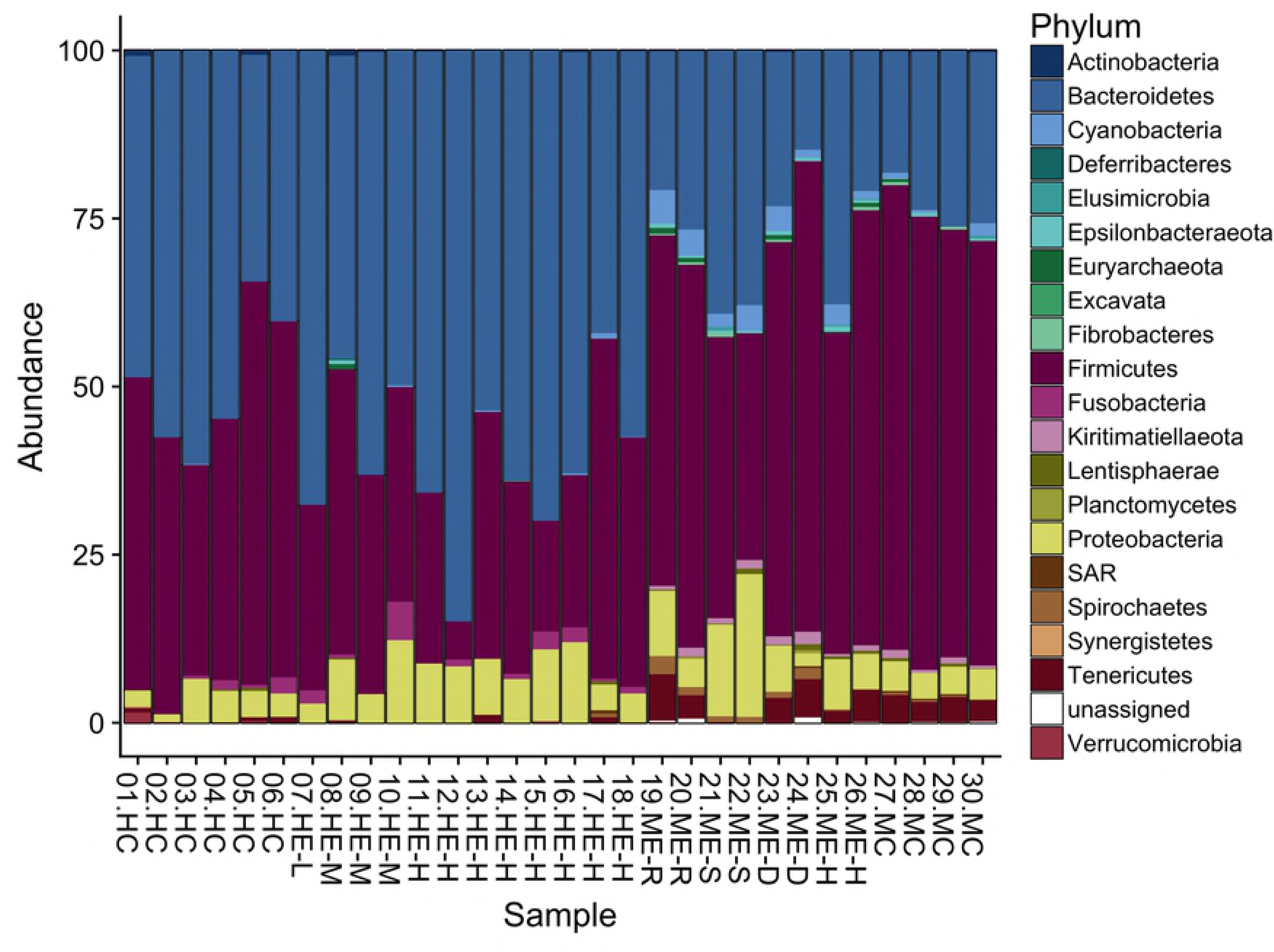
Phylum-level abundance bar plot. Relative abundance of bacterial phyla in macaque control (n=4), macaque exposed (n=8), human exposed (n=12), and human control (n=6) samples following rarefaction to the minimum library size. Samples are labelled with their record ID, group (HC, Human Control; HE, Human Exposed; ME, Macaque Exposed; MC, Macaque Control), and, if applicable, level of exposure (L, Low; M, Medium; H, High) and social group (R, S, D, H).

Using ANCOM analysis, no taxonomic features were significantly different in abundance between exposed and unexposed humans. Statistically significant differences in abundance between humans and macaques are listed in Table 5.

**Table 5.**
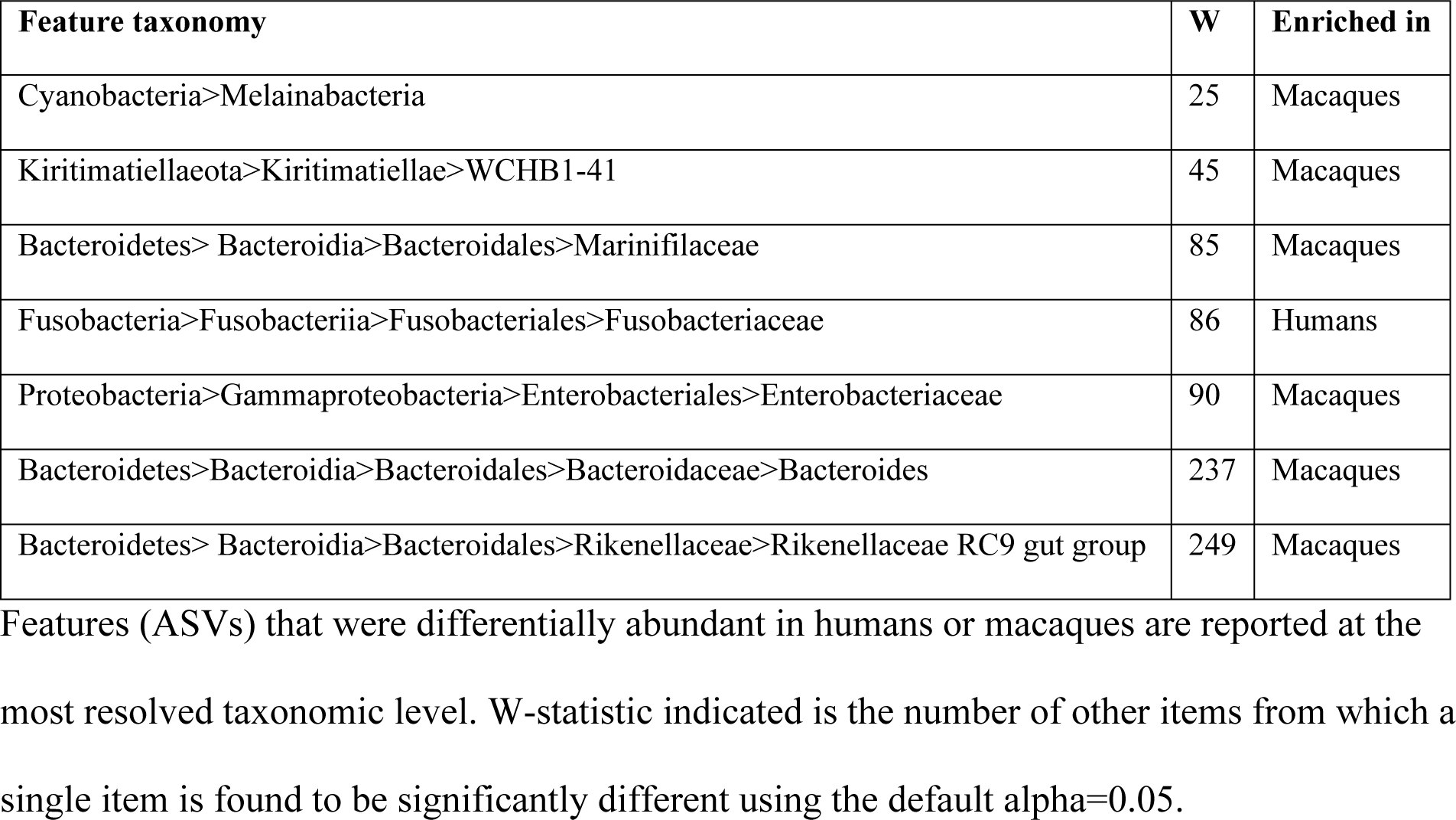
ANCOM analysis of differential abundance in humans and macaques

### Alpha and beta diversity

Fig. 2 displays the 1) total number of observed features in each sample, 2) Shannon’s index, which accounts for abundance and evenness of the taxa present using a natural logarithm, and 3) Simpson’s index, which measures the relative abundance of the different species making up the sample richness. For both humans and macaques, alpha diversity was consistently lower in the exposed groups (Fig 2), though this difference was only statistically significant among exposed humans compared to non-exposed controls.

**Figure 2.**
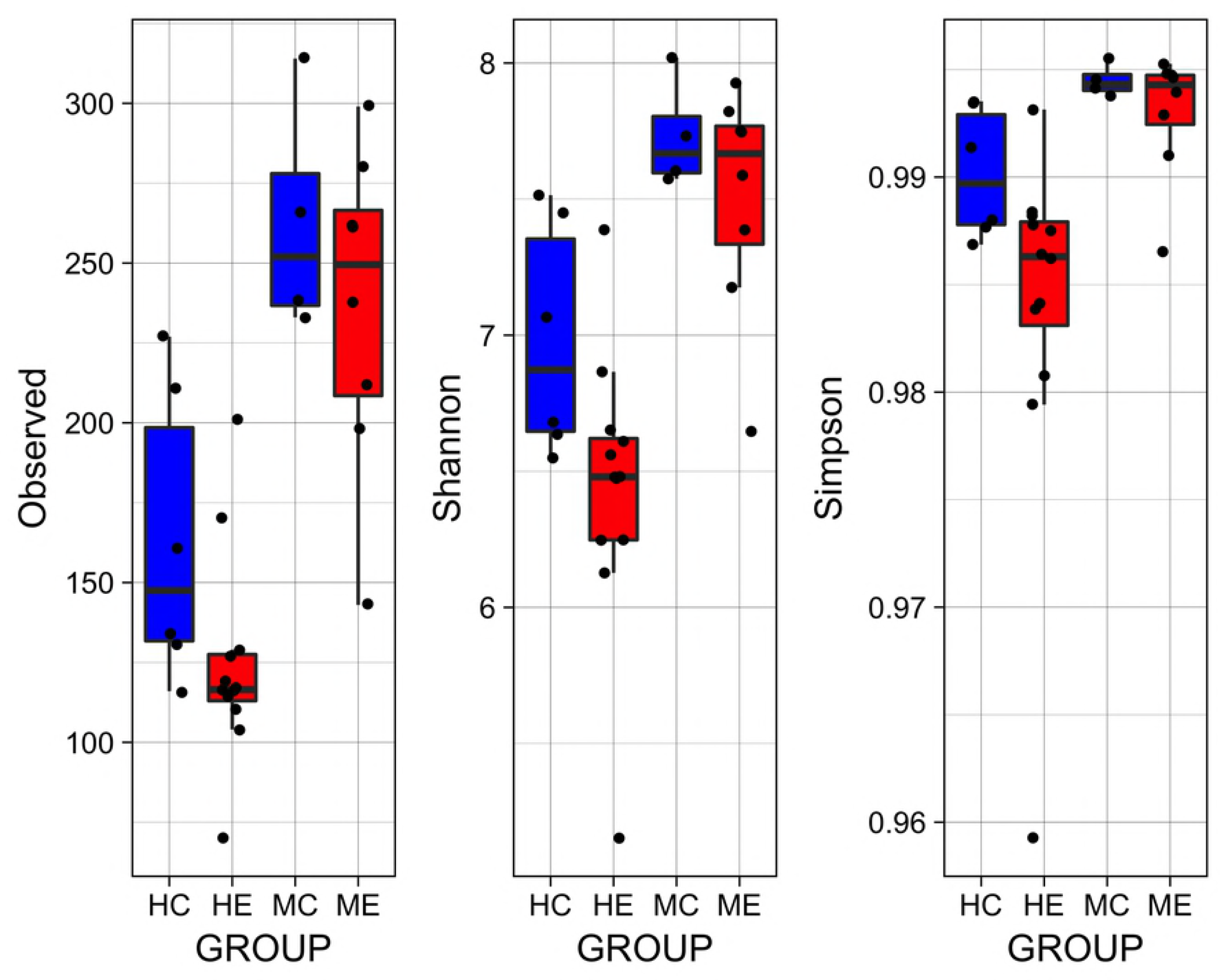
Alpha diversity. Alpha diversity was significantly lower among human exposed (HE) relative to human controls (HC) based on the observed features (p=0.04), Shannon’s index (p=0.02), and Simpson’s index (p=0.04). While there was a trend toward lower alpha diversity in exposed macaques, this was not statistically significant.

PCoA plots for human and macaque gut microbial communities are shown in Fig. 3. Dispersion using the betadisper test was significant for macaques (p=0.03), but not humans (p=0.66). Findings among humans were unchanged after excluding people who reported taking antibiotics in the past month (2 participants from control and 2 from exposed). Adonis [22] was used to test for location shift of the spatial median based on exposure status, and this statistic was significant for macaques and humans (p=0.04 and 0.02, respectively). Dispersion and location tests were also performed for unweighted UniFrac, Bray-Curtis, and Jaccard distance measures, with the same conclusions.

**Figure 3.**
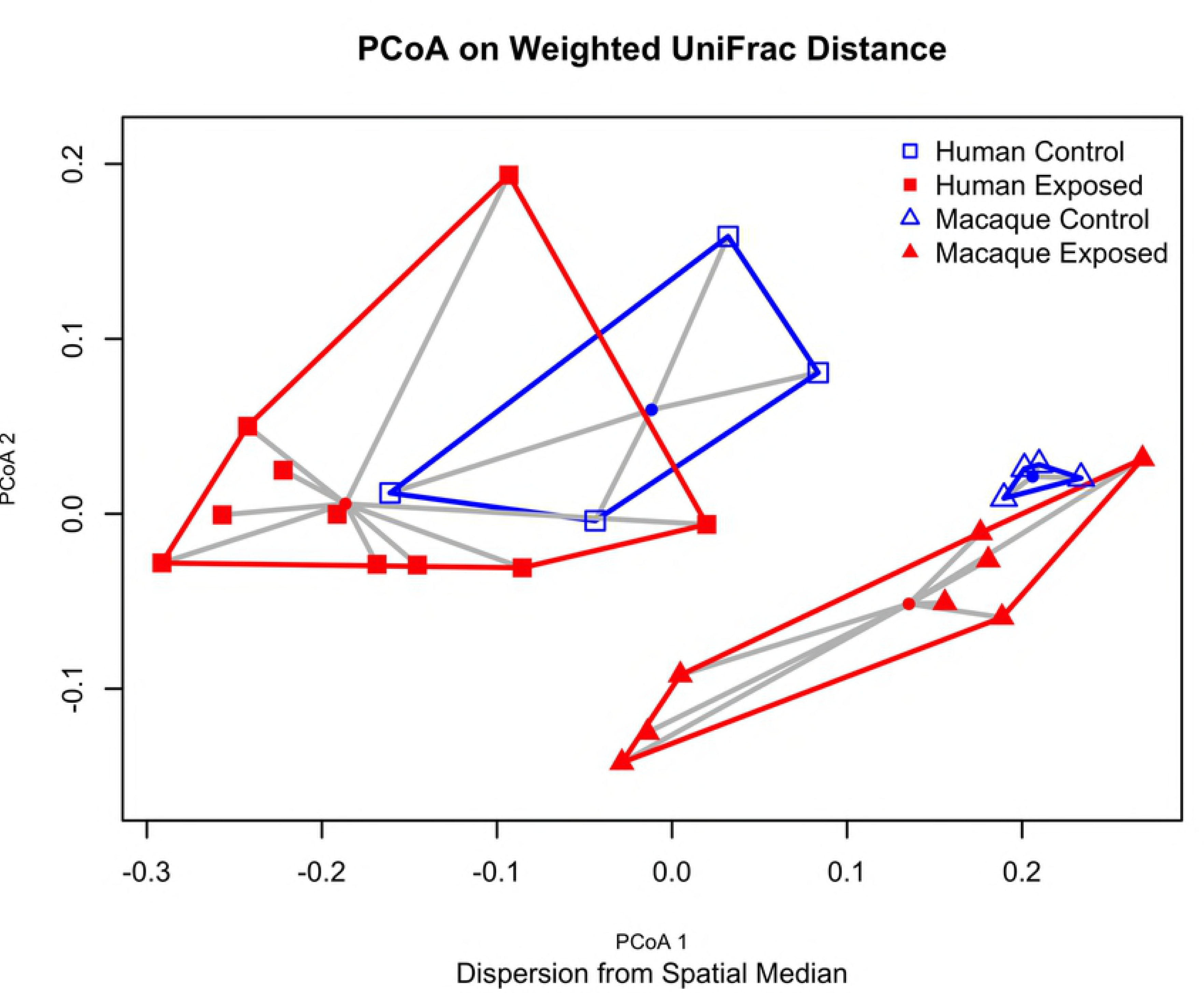
PCoA plot on weighted UniFrac distances. 2D PCoA plot based on weighted UniFrac distances demonstrate clustering and dispersion patterns for exposed human/macaque and control human/macaque samples.

### SourceTracker analysis

SourceTracker analysis (Fig. 4) revealed a higher percentage of microbes potentially sourced by macaque microbiota in the exposed human samples (mean=3.37%) compared to the controls (mean=1.84%). However, this difference was not significant by Mann-Whitney test (p=0.95) and was driven by higher proportions for one individual. Similarly, the reverse analysis, with humans as the source and macaques as the sink, showed a difference in proportions attributed to human samples for exposed and control macaques (mean=4.21% and 3.98%, respectively) that was not significant (p= 0.83).

**Figure 4.**
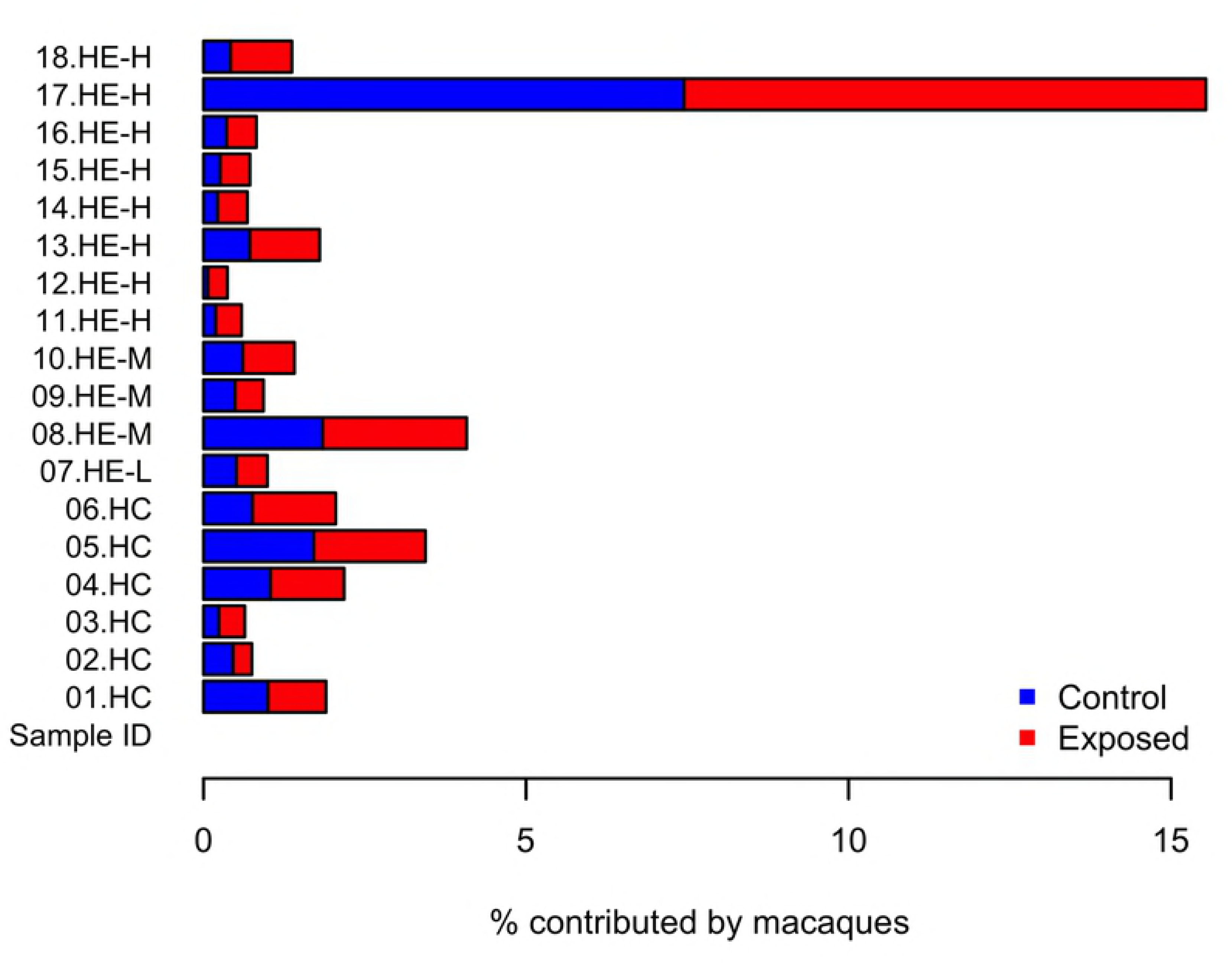
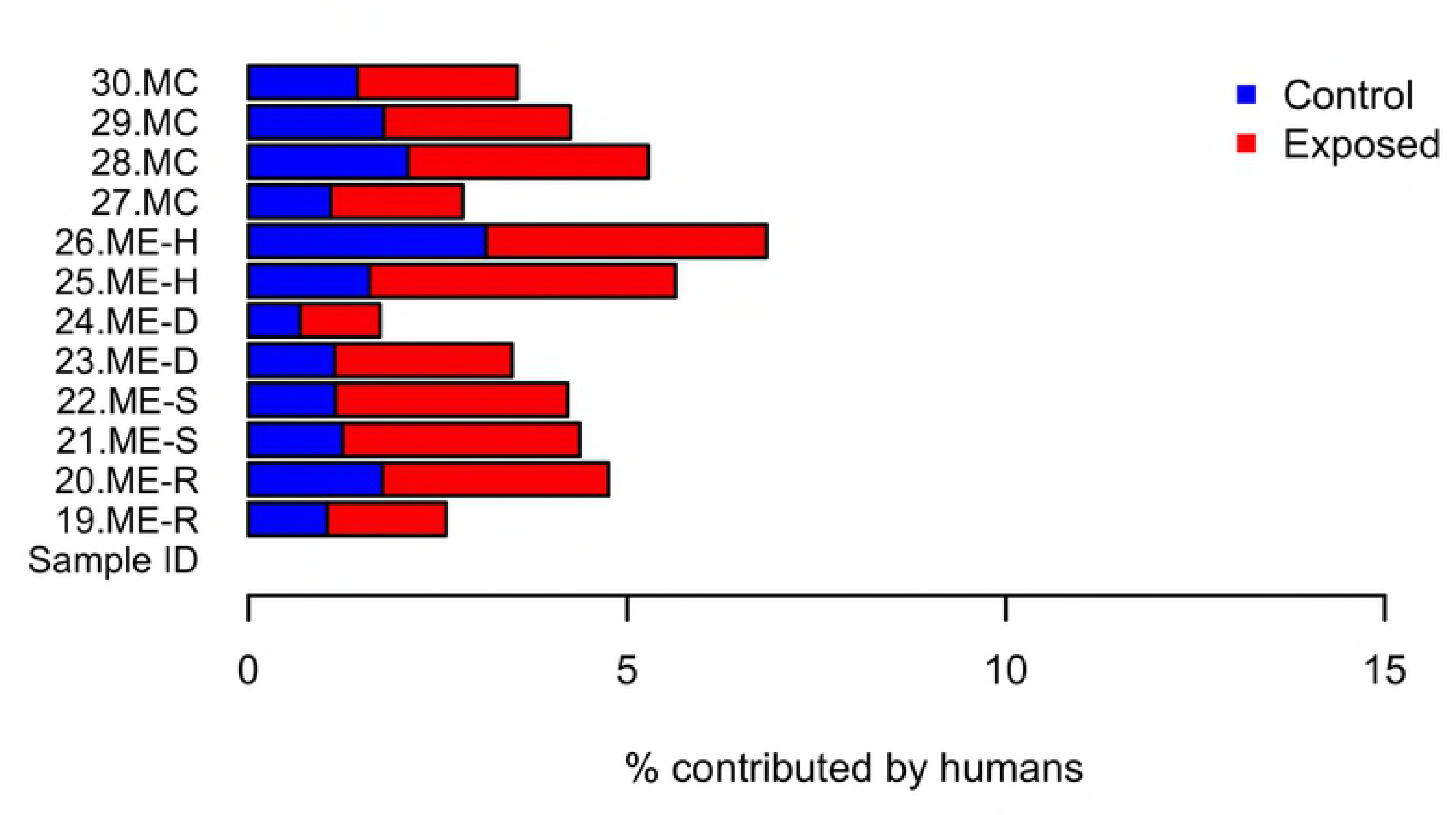
SourceTracker analysis. (A) Percent of each human sample attributed to macaque exposed or macaque control source, remainder is an unknown source. (B) Percent of each macaque sample attributed to human exposed or human control source, remainder is an unknown source.

## Discussion

Our study of gut microbiota in humans and macaques in close contact found that the degree of sharing between was not statistically significant. The gut microbiota of the exposed workers was significantly different from the control humans, although demographic differences could explain the shift. Exposed macaques in close contact with humans, compared to a less exposed population, exhibited beta-diversity dispersion effects that may reflect a dysbiotic, unstable gut microbiota composition, which may be tied human contact in an urban environment.

SourceTracker analysis revealed no significant difference in microbial sharing between humans and macaques However, one exposed worker had a greater proportion of their microbiota sourced from macaques than the other workers, suggesting that microbial sharing could be occurring and could depend largely on individual factors or behavior.

It is also worth noting that a common diet may play a role in the detected similarities, instead of or in addition to a shared environment. A study among urban Saudi and Bedouin populations compared to local baboons found that the shared environment and dietary overlap between Bedouins and local baboons resulted in more similar gut microbiome compositions relative to urban populations [25]. However, the relative importance of environment or diet was not characterized in that study. It is not clear to what degree the diet of macaques and humans overlap in our study setting. In addition to bananas, local residents and tourists bring a variety of foods to the macaques. During observations conducted between Sept-Dec 2016 [7] much of the provisioning consisted of fruits and vegetables, but also included chips, breads, and other foods not traditionally found in a macaque diet. Some macaques routinely foraged in trash or consumed more atypical food than others, so individual level dietary differences should be better characterized in the future.

Our analysis revealed that workers exhibited a different composition of fecal microbial communities than controls, as evidenced by significantly different spatial medians. This finding may be due to a number of other exposure factors that warrant further investigation to determine the consequences of this location effect, including differences in age, SES, smoking status, delivery mode, and history of infectious diseases. Early life factors are believed to play an important role in shaping the adult microbiome, and there were differences in delivery method and infant diet between exposed and control groups. While there is a considerable difference in age, all subjects were adults, so this factor alone is not expected to greatly influence results as gut microbiota, which tends to be well-established in healthy adults. Healthy adults’ gut microbiomes are usually less sensitive to perturbations than infants, whose microbiota are developing and have not reached a stable state and elderly (>75 years old), who tend to have lower total bacterial levels [26]. However, the difference age may be related to other factors (e.g., infectious disease history), which could shift their microbial composition. We also emphasize the need to exercise caution when excluding participants based on antibiotics use. Some respondents listed paracetamol or anthelminthic medication when asked about antibiotics use, or indicated they are unsure whether a drug they took was an antibiotic. Future studies should ask participants about antibiotic use by referencing specific drugs based on locally used names and example pills or obtain packaging from the medications used, if possible. A sub-analysis excluding participants who reported antibiotic use did not alter the general conclusions of this study.

The macaques in the park have a high level of gut microbiota dispersion relative to the macaques with minimal human contact. Dispersion essentially reflects variation of microbiota composition, that is the taxa present and their abundance differs from sample to sample among exposed macaques, whereas the control macaques are composed of similar taxa at a similar abundance, and therefore cluster tightly together, with minimal dispersion. This significant dispersion pattern on exposed macaques is suggestive of the “Anna Karenina principle,” a signature of dysbiosis characterized by increased variation in profiles of individuals in a disease state [27]. This dysbiosis may be due to environmental stressors or diseases that perturb a stable state in an unpredictable manner. We cannot definitively determine whether AKP effects are occurring without longitudinal sampling, however the initial findings are suggestive of these effects. In the KFP population, this dysbiosis could be a result of increased stress and competition among macaques, an increased disease burden, or may be attributable to their atypical diet. When asked if they noticed any changes in macaque behavior, workers reported that the macaques drank more Coca-Cola and ate more chicken than they used to. While most of the provisioned food appears to be fruits and vegetables, according to author RCK, who has observed this population extensively, the more extreme dietary changes like foraging in trash, might explain the high variation in composition among macaques at KFP. The population density of the macaques in KFP also is approximately 3,670 individuals/km^2^ which is considerably higher than found in more natural settings [7]. This likely results in elevated stress and aggression among macaques, which may ultimately facilitate pathogen spread. Since their microbiota appear to be in a dysbiotic state relative to macaques with low levels of human contact, a condition that may predispose them to gut-related diseases, they might be expected to present a greater health threat to humans than wild macaques with typical gut flora [28, 29].

Given that AKP effects are associated with growths of opportunistic pathogens, we expect to find lower evenness among the exposed macaques. While evenness based on Shannon index, Pielou evenness, and Simpson evenness was marginally lower among exposed macaques, this difference was not statistically significant. Our small sample size limited power and increased the risk of beta error, in which a study may fail to reject the null hypothesis due to insufficient power. This is a limitation that should be addressed in any future studies. By further characterizing changes in susceptibility to pathogens related to gut dysbiosis, we can improve understanding of the consequences of human activities such as diet supplementation or habitat encroachment on wild macaque populations.

We also found a location shift in the spatial medians of control and exposed macaque sample. It should be noted that, since the assumption of equal group variances is violated among macaques, the test used is not technically valid, however, since the group larger sample size is the same that exhibits greater dispersion, the test is liable to be too conservative, therefore the detected shift in spatial medians likely represents a statistically significant finding [30].

Another limitation of the study was the choice of human controls, who differed in many aspects from the exposed human population. As a result, microbiota differences between the groups could be due to demographic differences rather than factors related to macaque contact or occupation.

Further research should 1) investigate temporal trends and the stability of the dysbiosis described in this study, 2) recruit well matched controls (e.g. matched age, SES, gender) in Kosum Phisai to minimize the number of confounding factors in microbiota comparisons, and 3) incorporate testing for GI parasitism since many members of this community take anti-helminthic medication prophylactically and both factors have been shown to alter gut microbiota. The cross-sectional study design employed is practical as a baseline assessment that could be repeated in the future for continued, longitudinal surveillance of high risk worker populations and matched controls.

While the threat of acquiring an infectious disease shed through macaque feces from their work tasks appears low, we recommend that basic PPE be used, such as closed-toe shoes, to reduce the risk of acquiring environmentally transmitted parasites shed in macaque feces, which can enter through the skin. The high number of hand-to-mouth activities and work without respiratory protection represents a pathway for transmission of microbes that may be present in aerosolized macaque feces and offers insight into possible risk mitigating interventions. Due to the small sample size, we elected not to use scores from task observation videos in microbial composition analysis, instead treating all workers as exposed, however it is worth noting that the degree of exposure does indeed vary within this group. One worker noted that they experienced respiratory issues, which they attributed to the sweeping of macaque feces. Even if there were no microbial hazards from this exposure, the dust particles or endotoxin from Gram negative bacteria can cause irritation to the lungs. Therefore, future studies could assess lung function in the workers to help determine whether this workplace exposure contributes to decreased lung function or increased inflammation, respectively. Use of a mask during such tasks or misting of the ground prior to sweeping may reduce exposure to aerosolized macaque feces and protect worker health.

The use of a One Health approach, involving researchers from a range of disciplines, allowed us to compare the microbial status for both humans and the macaques of Kosumpee Forest Park. A traditional approach concerning only the human health impacts of macaque exposure may not have yielded an overtly apparent risk. Our finding of dysbiosis in the gut flora of macaques with close human contact may indicate increased susceptibility to pathogens. While further investigation is needed to determine the implications of this finding, a microbiome-based approach considering human and animal health in parallel may provide a more complete picture of health in an ecosystem.

## Conclusions

This study draws on a One Health approach to reduce human-animal conflict in a setting modified by habitat encroachment and fragmentation. Characterizing shifts in gut microbial communities allows for improved understanding of whether health changes are occurring due to increased human-macaque contact in a shared environment. Based on our analysis, workers’ gut microbiota may under certain circumstances be influenced by their exposure to macaques, but this would need to be confirmed in larger studies. The dispersion effects seen in macaques of Kosumpee Forest Park suggest that their altered diet and/or interaction with an urban environment may contribute to gut dysbiosis with unknown health consequences. Given the risk of transmission or respiratory irritation from ingesting or inhaling fecal microbes, exposed individuals and their employers should consider greater use of basic PPE and infection prevention methods, particularly proper hand hygiene. Targeted health protection and disease awareness promotion among forest park workers could limit opportunities for disease spillover from macaque populations into the broader community.

## Acknowledgements

We extend our gratitude to the study participants, Mr. Apichat Karaket, Director of Kosumpee Forest Park and park staff, and the staff of the Mattayom Watklangkosum School who helped facilitate this work. Many thanks to Janna Schurer and Gemina Garland-Lewis of the University of Washington Center for One Health Research for their guidance and support in manuscript revisions and IRB approval, respectively.

## Supporting information

**S1 File. Occupational exposure to macaques survey.** Questionnaire administered to workers to assess demographic, life history, diet, and general health, in addition to knowledge, attitudes and practices surrounding macaque exposure and zoonoses. Control surveys contained demographic, life history, diet, and general health sections only.

## References

1. Malaivijitnond S, Hamada Y. Current situation and status of long-tailed macaques (Macaca fascicularis) in Thailand. Nat Hist J Chulalongkorn Univ. 2008;8:185–204.

2. Kyes R, Jones-Engel L, Iskandar E, Onibala J, Lapin B, Chalise M, et al. Primate conservation biology in the 21st century: Global partnerships in research, training and outreach. American Journal of Primatology. 2006;68:144-. PubMed PMID: WOS:000239456400232.

3. Hankenson FC, Johnston NA, Weigler BJ, Di Giacomo RF. Zoonoses of occupational health importance in contemporary laboratory animal research. Comparative medicine. 2003;53(6):579–601.

4. Thanchomnang T, Intapan PM, Sanpool O, Rodpai R, Tourtip S, Yahom S, et al. First molecular identification and genetic diversity of Strongyloides stercoralis and Strongyloides fuelleborni in human communities having contact with long-tailed macaques in Thailand. Parasitology Research. 2017:1–7. doi: 10.1007/s00436-017-5469-z.

5. Wenz-Mücke A, Sithithaworn P, Petney TN, Taraschewski H. Human contact influences the foraging behaviour and parasite community in long-tailed macaques. Parasitology. 2013;140(6):709–18. doi: 10.1017/S003118201200203X. PubMed PMID: 23363557.

6. Gao F, Bailes E, Robertson DL, Chen Y, Rodenburg CM, Michael SF, et al. Origin of HIV-1 in the chimpanzee Pan troglodytes troglodytes. Nature. 1999;397(6718):436.

7. Kyes R, Tanee T, Thamsenanupap P, Karaket A, Kyes P. Population status of the long-tailed macaques (Macaca fascicularis) at Kosumpee Forest Park, Maha Sarakham, Thailand. American Journal of Primatology. 2017.

8. Village Health Volunteer Database Ministry of Public Health Thailand; 2014 [2017 May 19]. Available from: http://www.thaiphc.net/.

9. Ramirez V, Rabinowitz P, Kyes RC, Schurer JM, Grant ET, Trufan S, et al. Long-tailed macaques (Macaca fascicularis) in urban landscapes: Gastrointestinal parasitism and barriers for healthy co-existence in northeast Thailand. American Journal of Tropical Medicine & Hygiene. 2018;Manuscript submitted for publication.

10. Gregory AE, Lisa J-E, Michael S, Komang Gde S, Artha P, Agustin F, et al. Human Exposure to Herpesvirus B–Seropositive Macaques, Bali, Indonesia. Emerging Infectious Disease journal. 2002;8(8):789. doi: 10.3201/eid0808.010467.

11. Kim D, Hofstaedter C, Zhao C, Mattei L, Tanes C, Clarke E, et al. Optimizing methods and dodging pitfalls in microbiome research. Microbiome. 2017;5. doi: 10.1186/s40168-017-0267-5. PubMed PMID: WOS:000400644500002.

12. Song SJ, Lauber C, Costello EK, Lozupone CA, Humphrey G, Berg-Lyons D, et al. Cohabiting family members share microbiota with one another and with their dogs. elife. 2013;2.

13. Mosites E, Sammons M, Otiang E, Eng A, Noecker C, Manor O, et al. Microbiome sharing between children, livestock and household surfaces in western Kenya. Plos One. 2017;12(2). doi: 10.1371/journal.pone.0171017. PubMed PMID: WOS:000396161200073.

14. Harris PA, Taylor R, Thielke R, Payne J, Gonzalez N, Conde JG. Research electronic data capture (REDCap)—a metadata-driven methodology and workflow process for providing translational research informatics support. Journal of biomedical informatics. 2009;42(2):377–81.

15. Gilbert JA, Meyer F, Antonopoulos D, Balaji P, Brown CT, Brown CT, et al. Meeting report: the terabase metagenomics workshop and the vision of an Earth microbiome project. Standards in genomic sciences. 2010;3(3):243.

16. Caporaso JG, Kuczynski J, Stombaugh J, Bittinger K, Bushman FD, Costello EK, et al. QIIME allows analysis of high-throughput community sequencing data. Nature methods. 2010;7(5):335.

17. Callahan BJ, McMurdie PJ, Rosen MJ, Han AW, Johnson AJ, Holmes SP. DADA2: High-resolution sample inference from Illumina amplicon data. Nat Methods. 2016;13(7):581–3. Epub 2016/05/23. doi: 10.1038/nmeth.3869. PubMed PMID: 27214047; PubMed Central PMCID: PMCPMC4927377.

18. Weiss S, Xu ZZ, Peddada S, Amir A, Bittinger K, Gonzalez A, et al. Normalization and microbial differential abundance strategies depend upon data characteristics. Microbiome. 2017;5(1):27. Epub 2017/03/03. doi: 10.1186/s40168-017-0237-y. PubMed PMID: 28253908; PubMed Central PMCID: PMCPMC5335496.

19. Quast C, Pruesse E, Yilmaz P, Gerken J, Schweer T, Yarza P, et al. The SILVA ribosomal RNA gene database project: improved data processing and web-based tools. Nucleic Acids Res. 2013;41(Database issue):D590-6. Epub 2012/11/28. doi: 10.1093/nar/gks1219. PubMed PMID: 23193283; PubMed Central PMCID: PMCPMC3531112.

20. Mandal S, Van Treuren W, White RA, Eggesbø M, Knight R, Peddada SD. Analysis of composition of microbiomes: a novel method for studying microbial composition. Microb Ecol Health Dis. 2015;26:27663. Epub 2015/05/29. PubMed PMID: 26028277; PubMed Central PMCID: PMCPMC4450248.

21. McMurdie PJ, Holmes S. phyloseq: an R package for reproducible interactive analysis and graphics of microbiome census data. PLoS One. 2013;8(4):e61217. Epub 2013/04/22. doi: 10.1371/journal.pone.0061217. PubMed PMID: 23630581; PubMed Central PMCID: PMCPMC3632530.

22. Oksanen J, Kindt R, Legendre P, O’Hara B, Stevens MHH, Oksanen MJ, et al. The vegan package. Community ecology package. 2007;10:631–7.

23. Knights D, Kuczynski J, Charlson ES, Zaneveld J, Mozer MC, Collman RG, et al. Bayesian community-wide culture-independent microbial source tracking. Nat Methods. 2011;8(9):761–3. Epub 2011/07/17. doi: 10.1038/nmeth.1650. PubMed PMID: 21765408; PubMed Central PMCID: PMCPMC3791591.

24. Davis MF, Rankin SC, Schurer JM, Cole S, Conti L, Rabinowitz P, et al. Checklist for one health epidemiological reporting of evidence (COHERE). One Health. 2017;4:14–21.

25. Angelakis E, Yasir M, Bachar D, Azhar EI, Lagier J-C, Bibi F, et al. Gut microbiome and dietary patterns in different Saudi populations and monkeys. Scientific Reports. 2016;6:32191.

26. Lozupone CA, Stombaugh JI, Gordon JI, Jansson JK, Knight R. Diversity, stability and resilience of the human gut microbiota. Nature. 2012;489(7415):220–30. doi: 10.1038/nature11550. PubMed PMID: 22972295; PubMed Central PMCID: PMCPMC3577372.

27. Zaneveld JR, McMinds R, Thurber RV. Stress and stability: applying the Anna Karenina principle to animal microbiomes. Nature microbiology. 2017;2(9):17121.

28. Petersen C, Round JL. Defining dysbiosis and its influence on host immunity and disease. Cellular microbiology. 2014;16(7):1024–33.

29. Van Den Elsen LW, Poyntz HC, Weyrich LS, Young W, Forbes-Blom EE. Embracing the gut microbiota: the new frontier for inflammatory and infectious diseases. Clinical & translational immunology. 2017;6(1).

30. Anderson MJ, Walsh DC. PERMANOVA, ANOSIM, and the Mantel test in the face of heterogeneous dispersions: what null hypothesis are you testing? Ecological monographs. 2013;83(4):557–74.

